# *De novo* construction of a transcriptome for the stink bug crop pest *Chinavia impicticornis* during late development

**DOI:** 10.1101/2020.12.04.412270

**Authors:** Bruno C. Genevcius, Tatiana T. Torres

**Affiliations:** Department of Genetics and Evolutionary Biology, University of São Paulo, São Paulo, SP, Brazil

**Keywords:** genes, green stink bug, larvae, mRNA, nextGen, nymphs, sequencing

## Abstract

*Chinavia impicticornis* is a Neotropical stink-bug (Hemiptera: Pentatomidae) with economic importance for different crops. Little is known about the development of the species, as well as the genetic mechanisms that may favor the establishment of populations in cultivated plants. Here we conduct the first large-scale molecular study with *C. impicticornis*. We generated RNA-seq data for males and females, at two immature stages, for the genitalia separately and for the rest of the body. We assembled the transcriptome and conduct a functional annotation. *De novo* assembled transcriptome based on whole bodies and genitalia of males and females contained around 400,000 contigs with an average length of 688 bp. After pruning duplicated sequences and conducting a functional annotation, the final annotated transcriptome comprised 39,478 transcripts of which 12,665 had GO terms assigned. These novel datasets will provide invaluable data for the discovery of molecular processes related to morphogenesis and immature biology. We hope to contribute to the growing research on stink bug *evo-devo* as well as the development of bio-rational solutions for pest management.

## DATA DESCRIPTION

### Context

The stink bug *Chinavia impicticornis* (Hemiptera, Pentatomidae) is a Neotropical species with wide distribution across the South America. The species is an economically important polyphagous pest reported to feed on fourteen plant species [1]. Most of its damage has been attributed to soybean and closely related crops. However, the capability of switching to less preferred hosts in anomalous conditions may help explain the successful establishment of its populations in different crops [2,3].

Not only *C. impicticornis* but many other species of stink bugs are responsible for millions of dollars’ worth of damage worldwide every year. Taken as an example, the brown marmorated stink bug (*Halyomorpha halys*) has caused a 37 million dollar loss only to apple growers in the USA during 2011 [4]. Towards mitigating this damage, there is growing applied and basic research which has certainly contributed to our knowledge of the biological aspects and management strategies for stink bug pests [5]. Most efforts have focused on sampling methods, taxonomic identification, insecticide effectiveness, and population monitoring [6,7]. However, traditional pest control techniques have the potential of inflicting serious ecological disturbances as well as the selection of resistant lineages of crop pests. The development of bio-rational solutions is largely dependent on a detailed comprehension of the underlying biology of these pests [8]. In this context, studies on the “omics” fields offer promising species-specific and environmental-friendly tools [9]. More specifically, transcriptomic approaches provide the foundation for identifying gene targets associated with pheromones, pesticide resistance, and other features with potential for pest managing. Conversely, only five species of stink have had full body transcriptomes sequenced to date: the brown marmorated stink bug [10], the harlequin bug [9], the southern green stink bug [11], the brown stink bug [12], and the predatory stink bug *Arma chinensis* [13]. Genomic data are even more scarce, comprising four genomes assembled to date: *Halyomorpha halys* [14], *Euschistus heros, Piezodorus guildinii*, and *Stiretrus anchorago*.

Here we target this gap by documenting and characterizing the first transcriptome for the Neotropical stink bug *C. impicticornis* (Hemiptera, Pentatomidae). We conduct our study at the nymphal stage for two reasons. First, because management strategies are known to affect nymphs and adults in different ways, and nymph transcriptomes are yet scarce for stink bugs, *e*.*g*. [10]. Second, developmental transcriptomes provide invaluable data for the discovery of molecular processes underlying morphogenesis. We focused on the fifth instar because this is the stage where morphological sexual differentiation takes place [15]. These resources may be helpful for the growing research on genital *evo-devo* and stink bugs development as a whole [15,16].

## Methods

### Samples

Our colony of *C. impicticornis* was fed on green bean pods (*Phaseolus vulgaris*) and reared under the following conditions: 26±1 °C, 65±10% RH, and 14L:10D hours of photophase. We isolated RNA from male and female immatures at two developmental stages: the beginning and the end of the fifth nymphal instar (two hours after and seven days after molting from fourth to fifth instar, respectively). The fifth nymphal instar in our controlled conditions has an average of eight days. We included three individuals per sex per stage, totaling 12 specimens. We chose to remove the genitalia of each specimen to extract RNA and construct the libraries of bodies and genitalia separately. Genitalia are the most important traits for both species delimitation and sex identification in true bugs. Therefore, with this approach, we expect to generate data to be particularly useful for studying the mechanisms of sex determination and speciation. For RNA sequencing, each genital sample was sequenced separately (n = 3 genitalia per sex per sample), while bodies of the same sex and developmental stage were pooled to generate a single library (n = 1 body per sex per stage).

### RNA extraction & sequencing

Frozen specimens (at −80 °C) were sexed and had their genitalia separated under a stereoscope. RNA extraction of genitalia and bodies were conducted with the Trizol reagent following the manufacturer’s protocol (Invitrogen, Life Technologies). RNA was quantified via Qubit 2.0 (Thermo Fisher Scientific, USA) and then sent for sequencing at the Centro de Genômica Funcional (ESALQ-USP) using Illumina HiSeq 2500. Samples were paired-end sequenced with 100 bp and a target depth of 20 million read pairs per sample.

### Transcriptome assembly & annotation

We removed redundant sequences to decrease computational usage for transcriptome assembly using a custom Perl script [17] (supplementary material). *De novo* assembly was conducted in Trinity v. 2.4.0 [18] with concatenated samples, a minimum contig length of 199, and other parameters set to default values.

Due to the lack of a reference genome, transcriptome annotation was conducted using FunctionAnnotator [19]. This is a web-based tool that blasts sequences against the NCBI NR protein database. FunctionAnnotator was also used for the functional characterization, which uses the B2G4PIPE engine for GO terms assignment.

### Quality control

The quality of raw reads was assessed using the program FastQC v. 0.11.9. A quality control plot was constructed using MultiQC v. 1.9 [20] and is represented in Fig. 1. Raw reads of all samples had good quality. We removed low quality sequences and adapters using Trimmomatic v. 0.36 with the following parameters: LEADING:3 TRAILING:3 SLIDINGWINDOW:4:20 MINLEN:36. An average of 11.5 ± 0.006 % of raw reads was removed during the trimming process.

**Figure 1.**
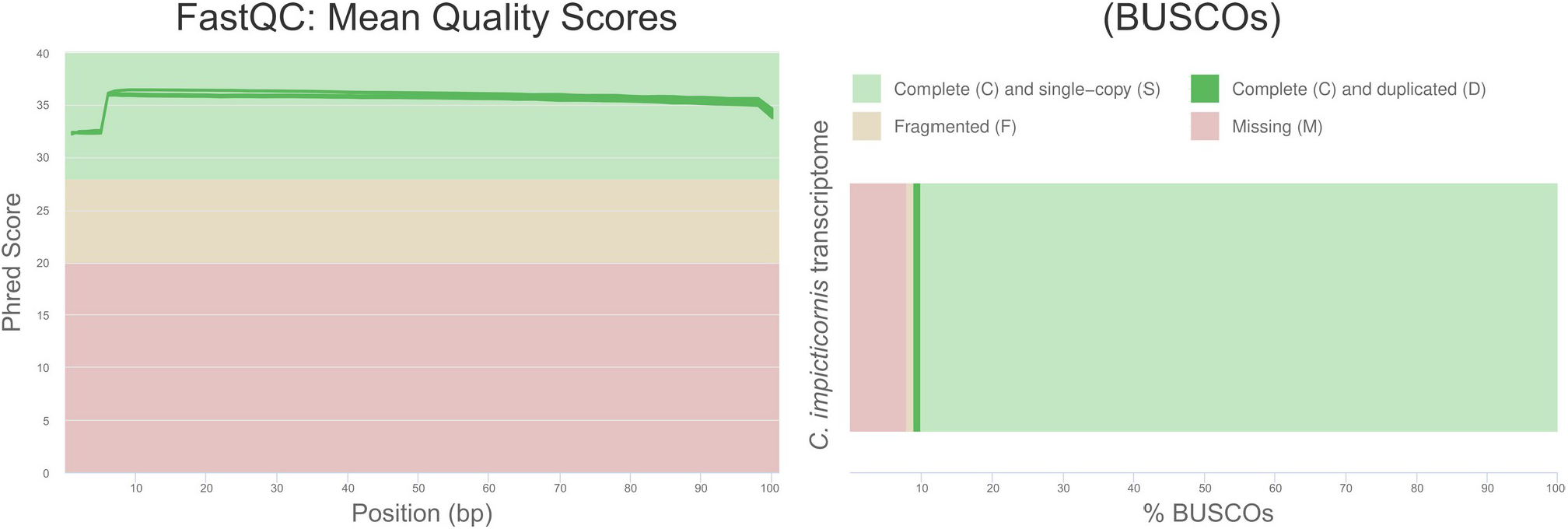
Quality control of raw reads using FastQC (left) and analysis of transcriptome completeness using BUSCO (right).

To avoid redundant transcripts in the transcriptome, we removed the shortest isoforms with a build-in Trinity script. We then assessed transcriptome completeness using the Benchmarking Universal Single-Copy Orthologs (BUSCO) v. 4.1.4 [21] against the Hemiptera ortholog database with default parameters. BUSCO analysis revealed appropriate levels of completeness for the assembled transcriptome (Complete: 90.1%, Fragmented: 1.5%, Missing: 8.4%; Fig. 1).

## Transcriptome characterization

RNA sequencing generated approximately 5 million raw reads (Table 1). The *de novo* assembled transcriptome had almost 400k contigs, with a 35.04% GC content and a mean contig length of 688 bp. After removing the shortest isoforms, 268,000 contigs remained. Out of these, 39,478 had successful matches in the NCBI database according to FunctionAnnotator, of which 36,329 aligned with metazoans and 33,871 with Arthropods. The vast majority of the contigs were annotated against the Hemiptera sequences, more specifically, against the genome of *Halyomorpha halys* (Fig. 2). All metrics had comparable values among previous studies with stink bugs (Table 1).

**Table 1.** Descriptive statistics of raw RNA-seq data and transcriptome assembly of the stink bug *C. impicticornis* compared with all other transcriptomic studies of pentatomids.

|  | *Chinavia*<br>*impicticornis* | *Arma*<br>*chinensis* | *Euschistus*<br>*heros* | *Halyomorpha*<br>*halys* | *Murgantia*<br>*histrionica* | *Nezara*<br>*viridula* |
| --- | --- | --- | --- | --- | --- | --- |
| Platform | Illumina<br>HiSeq. 2500 | Illumina<br>HiSeq. 2000 | Illumina<br>HiSeq. 2000 | Illumina<br>HiSeq. 1000 | Illumina<br>HiSeq. | Illumina<br>HiSeq. 2000 |
| Raw reads | 522,644,434 | nr | 126,455,838 | 439,615,225 | nr | 284,400,000 |
| Contigs<br>assembled | 392,739 | 112,029 | 83,114 | 248,569 | 526,403 | 299,148 |
| GC content | 35.04% | nr | 37.12% | nr | nr | nr |
| Mean contig<br>length | 688 bp | 250 bp | 1,000 bp | nr | 812 bp | nr |
| Pipeline | Trinity | Trinity | Trinity | Trinity | Trinity | Trinity |
| GOs<br>annotated | 5,087 | nr | 143,806 | 1,099 | 1,348 | 771 |
| Reference | This study | [13] | [12] | [10] | [9] | [11] |
| BioProject<br>accession | PRJNA666218 | PRJNA180995 | PRJNA488833 | PRJNA24284<br>9 | PRJNA302154 | PRJNA472074 |

**Figure 2.**
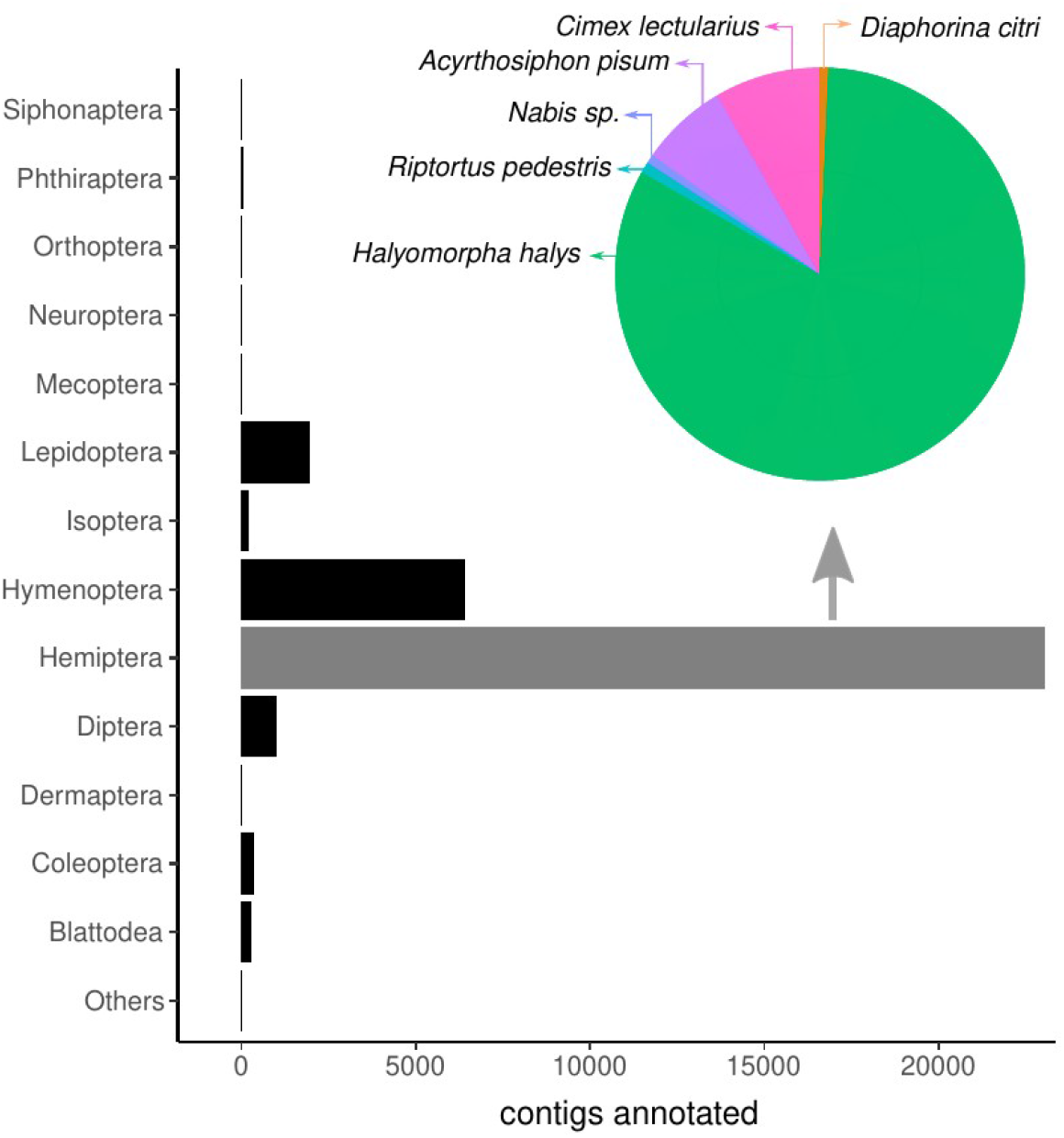
Diversity of insect orders from which the contigs of the *C. impicticornis* transcriptome were annotated.

For the 33,871 transcripts that aligned with arthropods, 12,665 transcripts had at least one GO term assigned. A total of 5,087 GO terms were identified combining the three categories: biological processes, molecular functions and cellular components. For the function “biological process”, the most represented GO term among the annotated transcripts was RNA-dependent DNA biosynthetic process (Fig. 3). For the molecular functions, RNA-binding was the most common GO term, while the cellular component with higher representativeness was the nucleus (Fig. 3).

**Figure 3.**
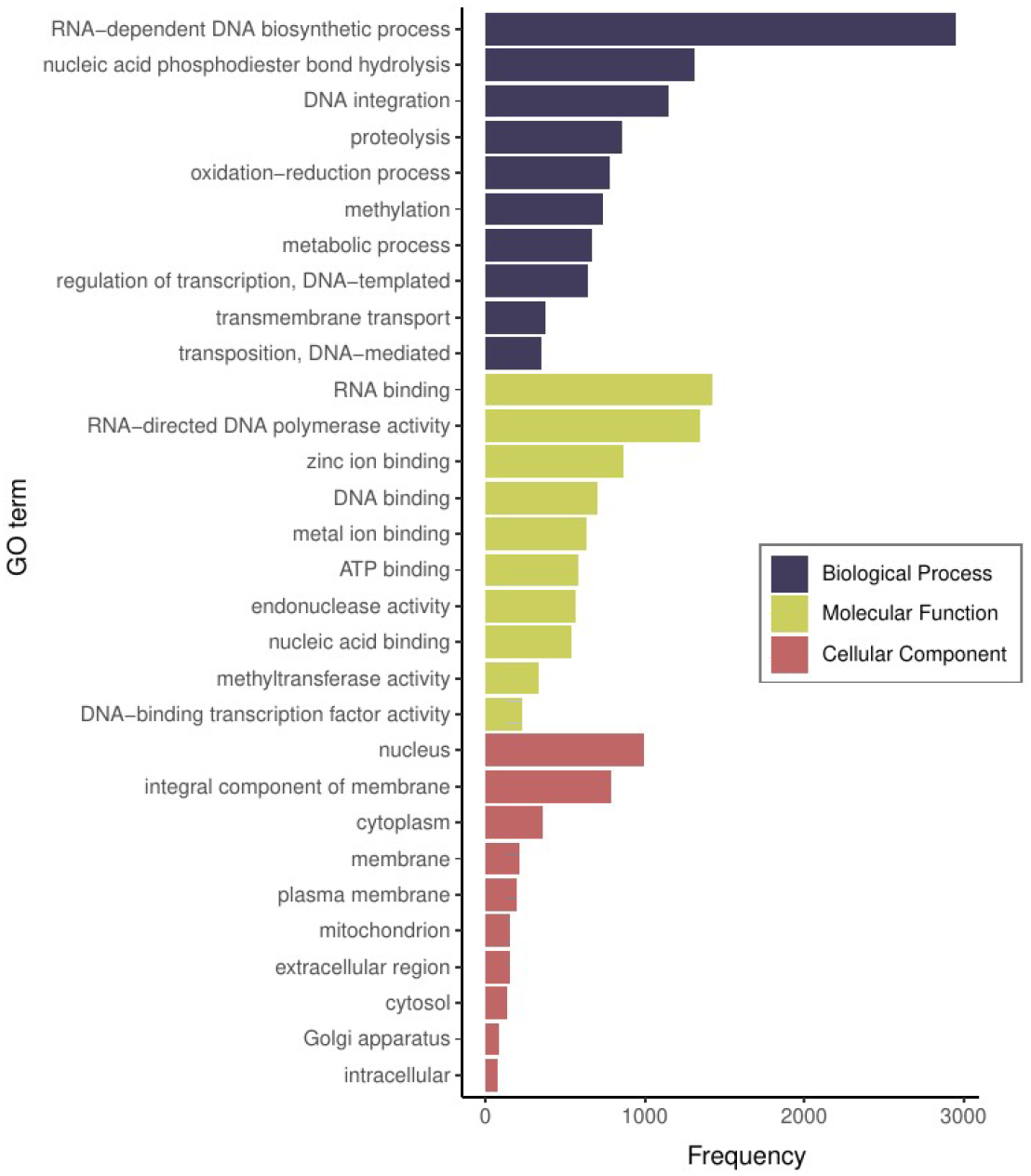
Top ten GO terms for each category with higher abundance in the transcriptome of *C. impicticornis* annotated with FunctionAnnotator.

## Re-use potential

Genomic and transcriptomic resources are paramount for advances in pest control and for the understanding of the origins of morphological innovations. Surprisingly, despite the high species diversity and the economic importance of stink bugs, large-scale comparative molecular studies are still scarce for this group of insects. Here, we present the first large-scale molecular study with the Neotropical pest *C. impicticornis*. From an economic perspective, we encourage further use of our data to determine the genetic bases of adaptation to new hosts and insecticide resistance. In particular, comparative transcriptomic studies may help explain why some stink bug species are more or less adaptable to different crops in comparison to *C. impicticornis*. From an evolutionary perspective, our data may be useful to investigate both the mechanisms of species separation and sexual differentiation. As genitalia tend to be the most complex and rapidly evolving traits in insects, studying the genetic/developmental architecture of these structures may help understand the emergence of morphological barriers during speciation. Our study is the first to make available a transcriptome of male and female immatures separately for stink bugs. We also provide, for the first time, transcriptome data of two different moments of a single immature stage, which will potentially inform about developmental heterochrony on a finer scale. Lastly, we also hope to contribute with useful data for phylogenomic studies of hemipterans.

## Availability of Supporting Data

Raw data are associated with the NCBI BioProject PRJNA666218 and are deposited at the Sequence Reads Archive (SRA) under the accession codes: SRR12762704, SRR12762703, SRR12762702, SRR12762696, SRR12762692, SRR12762689, SRR12762690, SRR12762691, SRR12762693, SRR12762694, SRR12762695, SRR12762697, SRR12762698, SRR12762699, SRR12762700 and SRR12762701. Transcriptome assembly is deposited at the NCBI Transcriptome Shotgun Assembly Sequence Database (TSA) under the accession code GIVF00000000.

## Acknowledgments

We thank Gisele Antoniazzi and Denis Callio for the assistance with establishing and maintaining the insect colony, as well as Luiza Saad and Federico Brown for their support with the RNA extractions. We also thank the reviewers Peter Thrope and Guillem Ylla and the editor Scott Edmunds for their constructive criticism during the review. BCG was supported by Fundação de Amparo à Pesquisa do Estado de São Paulo (FAPESP) with a Post-doctoral fellowship (proc. n. 18/18184-4). This work was supported by grants to TTT from FAPESP (grant 2016/09659-3).

## Authors’ contributions

BCG handled the insect colony, conducted the experiments and RNA extraction. BCG and TTT analyzed the data and wrote the manuscript.

## Competing interests

The authors declare no competing interests.

## Notes

https://www.ncbi.nlm.nih.gov/nuccore/GIVF00000000

### Competing Interest Statement

The authors have declared no competing interest.

